# Assessing the Effective Range for Individual Acoustic Identification: Comparison of Manual and Automatic Methods

**DOI:** 10.64898/2025.12.19.695494

**Authors:** Ilaria Morandi, Manuel Alejandro Jaramillo Rodriguez, Minkyung Kwak, Tereza Petrusková, Pavel Linhart

**Affiliations:** Department of Zoology, University of South Bohemia, České Budějovice, Czechia; Department of Electrical Engineering, KU Leuven, Leuven, Belgium; Department of Ecology, Charles University, Prague, Czechia; Department of Biology Ecoinformatics and Biodiversity, Center for Sustainable Landscapes Under Global Change, Aarhus University, Aarhus, Denmark

**Keywords:** Bioacoustics, Passive acoustic monitoring, Autonomous recording unit, Transmission test, Detection range, Population demography

## Abstract

Individual Acoustic Monitoring (IAM), especially when combined with passive acoustic monitoring (PAM), offers a non-invasive alternative to traditional mark-recapture methods to gain insights into species demography. Few studies have examined how identification decreases over distance along with degradation of identity cues. In this study, we conducted a song transmission experiment to quantify the range at which individual Yellowhammers (*Emberiza citrinella*) can be reliably identified. Songs from ten males (20 songs per individual, covering their full repertoire) were broadcast and re-recorded along a 200 m transect using AudioMoth recorders. Comparing manual classification by human observers with BirdNET classifier adapted for individual identification, we assess how well individuals could be distinguished at increasing distances. Humans were confident in assigning identity to a larger proportion of songs. Where identity has been assigned, both human and BirdNET were highly reliable at short distances (up to 50 m) discriminating even between very similar song types shared among males. At moderate distances (100 – 150 m), simple augmentation boosted BirdNETs performance remarkably, almost matching human classification accuracy. Our results indicate that individual recognition remains reliable up to 100 meters where both accuracy and agreement between assignment methods were high. Our results confirm that automated systems offer promising tools for large-scale, non-invasive individual monitoring, but challenges in accuracy and robustness persist at greater distances. We highlight the difference between signal detectability and individual identification range (much shorter) and its importance for optimizing PAM array designs.

## Introduction

Autonomous recording units (ARUs) have emerged as a scalable and increasingly popular technology for passive acoustic monitoring (PAM), providing cost-efficiency to traditional field observation methods (Shonfield and Bayne, 2017; Sugai and Llusia, 2019). ARUs are powerful tools for monitoring vocally active taxa (Gibb et al., 2019), with applications including species surveys, impact assessments of human activities, studies on climate change effects, and investigations into human–wildlife conflicts (Blumstein et al., 2011; Penar et al., 2020; Bota et al., 2022; Ross et al., 2023). In PAM studies, ARUs are typically deployed in the field and programmed to record over predefined periods, after which the collected audio data are analyzed. Recordings can be either listened to or visually inspected, although this process becomes increasingly challenging when the total recording time extends to several hundreds or thousands of hours (Towsey et al., 2018).

Many bird species produce clear and consistent vocalizations, thus opening the door for using automated signal recognition software to efficiently process large PAM acoustic datasets (Knight et al., 2024). The rapid development of machine learning and deep learning techniques has greatly enhanced the processing and analysis of extensive bioacoustics data. Neural networks including deep learning frameworks are now widely applied to automate acoustic detection and recognition tasks and handle large datasets (Stowell et al., 2019; Kahl et al., 2021; Denton et al., 2022; Stowell, 2022; Knight et al., 2024). User-friendly and ready-to-use machine learning tools have recently been developed and are increasingly accessible to respond to real-life monitoring challenges and the general public (Cole et al., 2022). Among these approaches, BirdNET (Kahl et al., 2021) and Google-Perch (Ghani et al., 2023) are two CNN-based models widely used for species classification (Ghani et al., 2023; Funosas et al., 2024). In particular, BirdNET Analyzer provides an easy-to-use graphical user interface (GUI), allowing users to upload audio recordings directly. By processing three-second audio segments, the algorithm of the latest BirdNET version (v2.3) is capable of identifying more than 6,000 species worldwide (Kahl et al., 2021). To date, however, most bioacoustics research employing such approaches has focused on species-level monitoring.

Currently, there is a growing interest in acoustic individual identification (AIID), which aims to move beyond species-level classification to identify individual animals through their unique vocal signatures (see, for example, the comprehensive reviews by Knight et al., 2024). AIID offers a novel approach to ecological and evolutionary research and monitoring, providing an alternative to mark–recapture techniques (Terry and McGregor, 2002). Case studies using focal recordings of individual birds have shown that AIID has the potential to track individuals across space and time. It can be used for censuses (e.g. Hensel et al., 2022), to estimate population size (e.g. Marin-Cudraz et al., 2019), to study dispersal (e.g. Odom et al., 2013) or site fidelity (e.g. Bailey et al., 2021), and has been shown to improve estimated survival rates relative to labor-intensive traditional methods (e.g. Vögeli et al., 2008). Indeed, the potential applications of AIID are numerous.

Most sound-producing taxa have acoustic signatures (Linhart et al., 2022; Knight et al., 2024; Madhavan and Linhart, 2025), but AIID has only been applied for a few of them usually requiring extensive manual analysis (e.g. Petrusková et al., 2016; Clink et al., 2017). Huang et al. (2025) demonstrated that models pretrained on bird species classification can be adapted and fine-tuned to classify vocalizations of known individual birds and detect unknown individuals. Specifically, they tested pretrained models such as AemNet (Lopez-Meyer et al., 2021), BirdNET, and Google-Perch, originally trained for species-level classification, and showed that these models can be effectively retrained for individual-level recognition. The models were evaluated under both closed- and open-set scenarios, providing strong support for the usefulness of pretrained models in individual acoustic recognition tasks. These findings indicate that automatic AIID approaches are feasible for species exhibiting robust and consistent individual acoustic signatures.

One of the major challenges in AIID is the lack of suitable datasets labeled with individual identities, preferably recorded under variable and acoustic conditions, which are needed to train deep learning models (Stowell et al., 2019). Most previous studies have relied on targeted focal recordings of known individuals collected with directional handheld recorders, which offer high signal-to-noise ratios but are often constrained by limited sample sizes (reviewed in Knight et al., 2024). The growing adoption of PAM and ARUs now allows the collection of large-scale, long-term bioacoustics datasets. However, identifying individuals from PAM recordings is challenging due to factors such as variation in recording conditions, signal degradation over distance and overlapping calls (Knight et al., 2024). Signal attenuation and degradation due to environmental factors (such as wind, vegetation density, etc.) have consequences for the ‘active space’ of bird sounds. Active space refers to the area around the sound source over which the signal remains detectable and recognizable (Brenowitz, 1982a). Also, a low signal-to-noise ratio will always result when the individual is positioned far away from the microphone, potentially leading to incorrect or uncertain individual assignment.

Assessing the active space of the various types of information encoded in songbirds’ vocalizations is critical for addressing ecological questions, such as individual spacing, and for understanding social behaviors, including territorial and mating strategies. While previous studies have primarily focused on the degradation of species-specific information (e.g., species identity (see Hardt and Benedict, 2021; Winiarska et al., 2024), there is a substantial knowledge gap regarding the transmission of finer-grained information, such as individual identity, through the environment.

Understanding the range over which signals remain detectable is essential for establishing robust protocols for PAM. A well-established practice to study signal active space and signal degradation is to broadcast recorded animal signals and record them at multiple distances, thereby simulating communication within and across animal territories (Hardt and Benedict, 2021). However, only a few studies have employed autonomous recording units (ARUs) to conduct signal transmission tests. These studies aimed to improve understanding of detection distances, which are species-specific and influenced by habitat and environmental characteristics (Yip et al., 2017; Priyadarshani et al., 2018; Winiarska et al., 2024). While there are a few studies that studied degradation of features allowing individual identification (e.g., White-browed Warbler: Mathevon et al., 2008; Corncrakes: Rek and Osiejuk, 2011; Zebra Finch: Mouterde et al., 2014), no study has defined the range of individual identification using ARUs and automatic processing pipelines that are required in PAM.

Numerous features make the Yellowhammer (*Emberiza citrinella*) a particularly suitable model species for studying AIID. It’s a common European bird, widespread across Eurasia, with a relatively simple song. Males can be recognized individually based on the types of songs they sing during the long breeding season (Hiett and Catchpole, 1982; Hansen, 1999), and they maintain a stable repertoire throughout their lives (Hansen and Balsby, 2012). Yellowhammers and their songs have attracted the attention of naturalists and scientists for over 150 years (e.g. Oppel, 1869) and they have long been a popular model species, particularly for studying dialects in birdsong (Petruskovà et al., 2015). Additionally, Yellowhammers serve as indicator of healthy farmland (Bradbury et al., 2000), with their populations declining rapidly across Europe (BirdLife International, 2024). The species’ individually distinctive songs may therefore provide a valuable, noninvasive tool to monitor rapid population changes.

In this study, we conducted a transmission experiment using ARUs (AudioMoth) to assess the range at which individual Yellowhammers can be reliably identified under PAM survey context. We combined manual (human) and automatic (ready-to-use deep learning) identification approaches to assign individual identity from recordings collected at increasing distances. We provide a valuable annotated dataset of individually labeled songs that can be used for further development of automatic detection tools, and along with provided human performance, it can be used as a benchmark for AI assisted AIID tasks.

## Materials and methods

### Subject

The Yellowhammer (*Emberiza citrinella*) is a common Palearctic passerine, with a wide distribution ranging from Spain to central Asia (Cramp and Perrins, 1994, del Hoyo et al., 2011). This species produces a short and stereotyped song composed of two distinct parts: an initial phrase and a dialect phrase (Figure 1a). The initial phrase has a trill-like structure, usually consisting of a rapid repetition of a pair of elements (Frauendorf, 2005), and shows high inter-individual variability (e.g. Hansen, 1984; 1999; Caro et al., 2009); Each male possesses a repertoire of 1 to 5 (mostly 2) initial phrase types (i.e. song variants) which remain stable throughout life (Hansen and Balsby, 2012). Individuals repeat these song variants in a stereotyped sequence to form a song bout before switching to a new song variant (Hansen, 1978; Rutkowska-Guz and Osiejuk, 2004). The combination of initial phrase types in the repertoire mostly differed between individuals and even when, in rare cases, males have identical repertoires, they could always be distinguished by minor differences in the song elements (Hansen and Balsby, 2012). In contrast, the dialect phrase, composed of 2 (occasionally 3) longer elements, reflects regional dialects (Petrusková et al., 2015; Diblíková et al., 2023) and provides population-level information.

**Figure 1.**
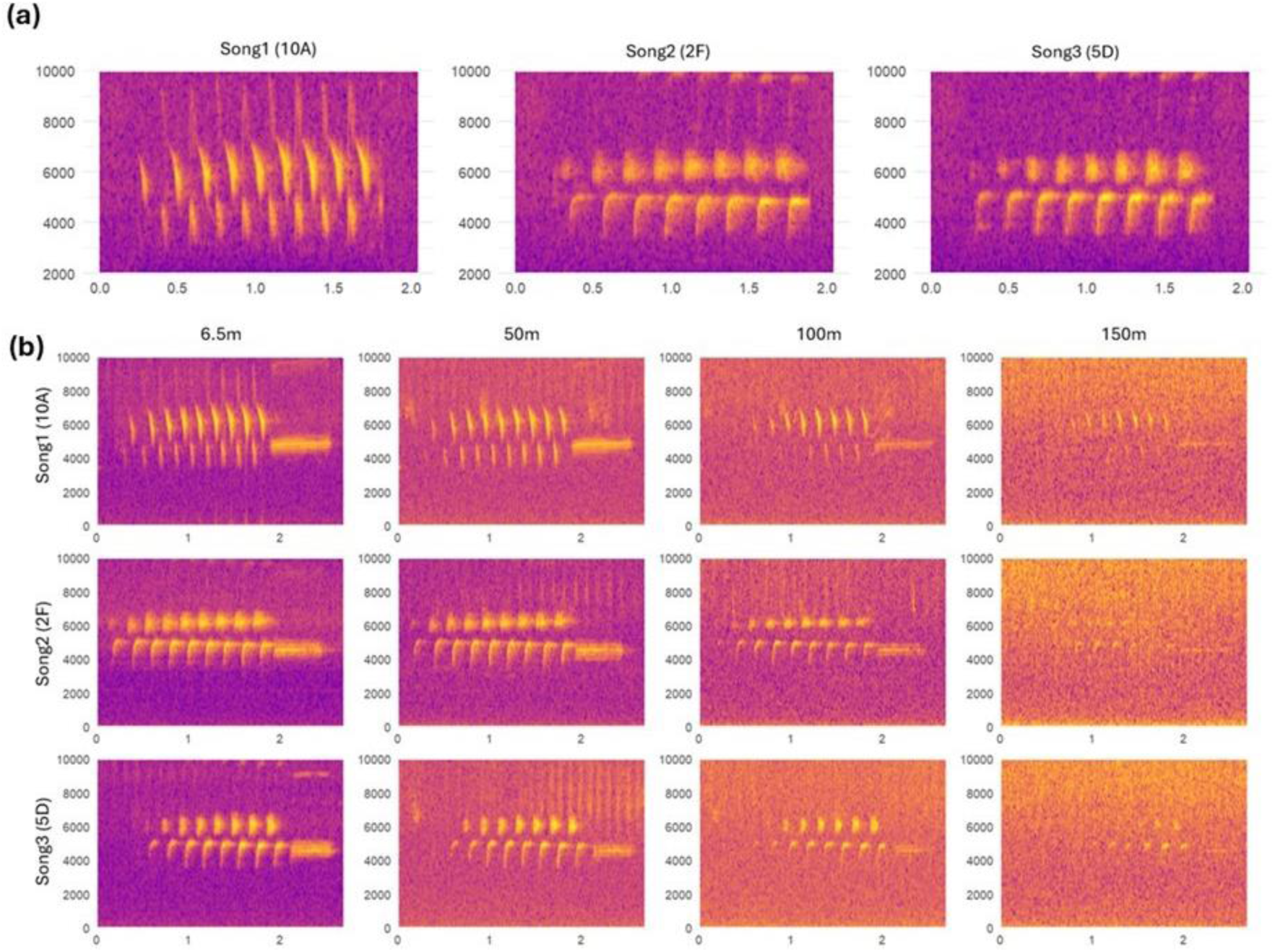
Spectrogram representations of Yellowhammer (*Emberiza citrinella*) songs. (a) High similarity is evident when males share their first phrases probably due to song imitation (Song2 and Song3) (see details in 2.4.1 Song prototypes). (b) Rows illustrate inter-individual variation while columns illustrate the effect of distance. Spectrograms plotted using the seewave package in R (Sueur et al., 2008).

### Propagated sounds

Yellowhammer songs were initially recorded with AudioMoth recorders placed near the birds’ favorite singing posts, less than 5 m from the singing birds. The series of songs was chosen from a larger set of recordings based on their quality. We selected the highest quality songs with high signal-to-noise ratio (assessed visually based on spectrograms), and no overlap from other conspecific or heterospecific songs. Using Audacity (v.3.7.1), we applied noise reduction on some songs (20 dB; sensitivity = 6.0, 3; frequency smoothing bands). Finally, we filtered (high-pass band filter; roll-off: 12 dB; 2500 or 2800 Hz; Yellowhammer song frequency range: approximately 2000 to 8000 Hz) and normalized each song from a single recording sequence (peak amplitude to -1.0 dB). Using these prepared sounds, we created a playback track by pasting the stimuli into a single file. The track contained 209 songs from 10 individuals, including in total 21 different initial phrase types (2-3 types per male), with c.a. 10 repetitions of each initial phrase type. It included an artificial beep at the outset (to mark the start and the end of the experiment) and each of the songs in succession, with 1.5 s of silence between each sound. This interval both prevented overlaps with the preceding song and subsequently allowed measurement of the signal-to-noise ratio for each song playback.

### Transmission experiments - Data collection

The sound propagation experiment was conducted on 26 July 2024 in Vižina (49°51′26″N, 14°6′17″E), located in the Central Bohemian Region of the Czech Republic. It was carried out on grassland, an open habitat consisting mainly of maintained and cultivated meadows with scattered shrubs and small trees. This habitat was chosen because it represents the typical breeding and singing environments of the Yellowhammer, a species strongly associated with semi-open and agricultural landscapes. We played back the Yellowhammer track from a loudspeaker (Vonyx ST014) mounted on a post at a height of 1.5 m. We broadcasted all the signals at 85 ± 3 dB SPL measured at 1 m from the loudspeaker with a sonometer (Voltcraft SL-200). This value corresponds to the natural amplitude of a Yellowhammer (Brackenbury, 1979). We re-recorded sounds in the morning between 0800 h and 0900 h with seven Audiomoths at different distances (6.50 m, 12.50 m, 25.0 m, 50.0 m, 100.0 m, 150.0 m, and 200.0 m). At each distance, we broadcasted the set of sound stimuli two times, in an attempt to maximize the chance that the stimuli would be recorded at least once without being overlapped by other sound sources in the field.

### Testing dataset preparation

We visually examined the sound files from each Audiomoth to determine if the re-recorded vocalizations could be seen on the spectrogram in each recording. If so, the detections were marked and assigned to the corresponding individual in the track. This was done by one person (IM) with prior knowledge of the order of male vocalizations in the original recording using Praat v. 6.4.23 (View range: 0 – 10,000 (Hz); Window length: 0.01 (s); Dynamic range: 50 (dB)). Dynamic range was occasionally adjusted to analyze recordings with lower signal-to-noise ratio (SNR). For each song at each recording distance, SNR was calculated, representing the relationship between the signal’s energy and background noise, using the “baRulho” package in R (Araya-Salas et al., 2025). Background noise was sampled as a 1-second time slice immediately preceding each song (Dabelsteen et al., 1993). The 1463 audio fragments containing the songs were then cut from the original recordings, and organized into folders, each corresponding to a specific distance. The files were then anonymized and randomized for the assignment task (testing dataset).

### Song prototypes

For individual identification of Yellowhammers, we created a catalog of the 10 males used in the playback experiment (Supporting information, Figure S1). This catalog included high quality spectrograms of their songs, each with an assigned code. Each prototype was then defined as a combination of a broader song phrase category (A-H) and an individual variant of that song phrase corresponding to the male ID (1-10). For example, prototype 1A is a variant of song phrase A sung by male number 1. Some prototypes were quite unique, while others were more similar. High similarity was particularly evident when males appeared to share the same song phrase. Yet, small consistent differences made it possible to distinguish between them (e.g. the frequency separation between alternating high and low frequency syllable elements in Song2 (2F) is larger than in Song3 (5D), see Figure 1a).

### Manual Individual Identification

The 1463 anonymous files (testing dataset) were opened and analyzed in Praat (v. 6.4.23) by two external observers (PL and MK). The observers used the same parameters as those applied by the initial observer with prior knowledge (View range: 0 – 10,000 (Hz); Window length: 0.01 (s); Dynamic range: 50 (dB)) occasionally adjusted slightly. The presence or absence of Yellowhammer songs was determined from the spectrograms, and each song was assigned to a particular individual following the prototype reported in the provided catalog (Supporting information, Figure S1). Prototype was not assigned in cases when observers judged the song to be too degraded and missing individually distinct traits.

### Automated Individual Identification

We used BirdNET-Analyzer (GUI version 1.5.0; BirdNET model version V2.4) to train two multiclass models discriminating between 21 different classes representing different song prototypes. Each prototype was included in the model as a distinct class and trained using 20 example songs of each prototype (420 songs in total). We selected these songs from the same song recordings but different songs that were used to create the transmission playback tape.

The first model, which we will call “BirdNET Basic”, was trained utilizing the default settings on BirdNET-Analyzer, with the exception of the minimum and maximum bandpass frequencies, which were adjusted to align with the frequency range of Yellowhammer songs (2000 and 9000 Hz, respectively). The remaining settings employed for this model are listed in Table 1. Our intention was not to create the optimal model, but rather to establish a baseline performance for the off-the-shelf classifier that has been successfully employed in numerous other contexts.

**Table 1:** Parameters used for data augmentation. We include the probability, minimum, and maximum values of the time shift, frequency shift and time stretch, as well as the target SNR values for each of the new generated samples.

|  | *Min Value* | *Max Value* | *Probability* |
| --- | --- | --- | --- |
| *Time Shift (seconds)* | -0.3 | 0.3 | 0.5 |
| *Frequency Shift (Tones)* | -0.2 | 0.2 | 0.5 |
| *Time Stretch (Ratio)* | 0.9 | 1.1 | 0.5 |
| *SNR target values* | [-15, -10, -5, 0, 5, 10, 15, 20, 25, 30] |  |  |

For the second model, “BirdNET Advanced”, we conducted a more sophisticated training procedure to achieve higher performance with the same available data. First, we used data augmentation, increasing the number of training songs per prototype. We created new synthesized recordings by applying four different types of modifications: (i) adding background noise, (ii) time stretching, (iii) pitch shifting and (iv) time shifting (Kahl et al., 2021). For the background noise augmentation, we set different target signal-to-noise ratios (SNRs) between the original song and the added noise; one new sample was created for each target SNR. The background noise samples were obtained from intervals between Yellowhammer songs in the same recordings from which the prototype songs were extracted. Finally, the other modifications (time stretch, pitch shift, and time shift) were randomly added according to the parameters ranges and probabilities reported in Table 1.

Second, we performed hyperparameter tuning with validation-based model selection, while maintaining the same model architecture provided by BirdNET-Analyzer. Specifically, we used a grid search approach to vary the number of hidden units and the up sampling ratio. The other hyperparameters were set to default. To perform validation, we applied the hyperparameter grid search for each of the augmented datasets generated by using different augmentation factors (x2, x3, x4, x6, x10). Then, we selected the best-performing model based on the validation metrics (Auroc and Auprc) reported by BirdNET-Analyzer. The hyperparameters of the selected model are reported in Table 1 and the target SNR values for the data augmentation in Table 2.

**Table 2:** Hyperparameters used during BirdNET basic and BirdNET advanced fine-tuning.

|  | *Epochs* | *Hidden units* | *Dropout* | *Batch Size* | *Learning rate* | *Upsampling mode* | *Upsampling ratio* | *Mix up* |
| --- | --- | --- | --- | --- | --- | --- | --- | --- |
| *BirdNET Basic* | 50 | 0 | 0.0 | 32 | 0.001 | Repeat | 0.0 | FALSE |
| *BirdNET Advanced* | 100 | 516 | 0.33 | 32 | 0.0001 | Repeat | 0.5 | TRUE |

### BirdNET confidence threshold

BirdNET predictions are accompanied by a quantitative confidence score ranging from 0 to 1, which indicates the model’s certainty that a given prediction has been correctly identified (Wood and Kahl, 2024). We set a confidence score threshold to filter BirdNET output and include only predictions exceeding a specified probability of being correct (e.g. Bota et al., 2023; Singer et al., 2024). We determined the optimal threshold balancing high precision and maximizing the number of predictions retained in the dataset (Tseng et al., 2025). We analyzed the overall performance of the main metric and calculated the number of songs retained at each confidence score for each model. Precision and the number of retained songs are shown in Figure 2, while the other metrics (accuracy, recall, and F1 score) are reported in Supporting information, Table S1. We selected a confidence threshold of 0.5 which represents a balance between model performance (0.9 for BirdNET Advanced) and the number of songs retained (63% for BirdNET Advanced), avoiding the exclusion of too many examples.

**Figure 2.**
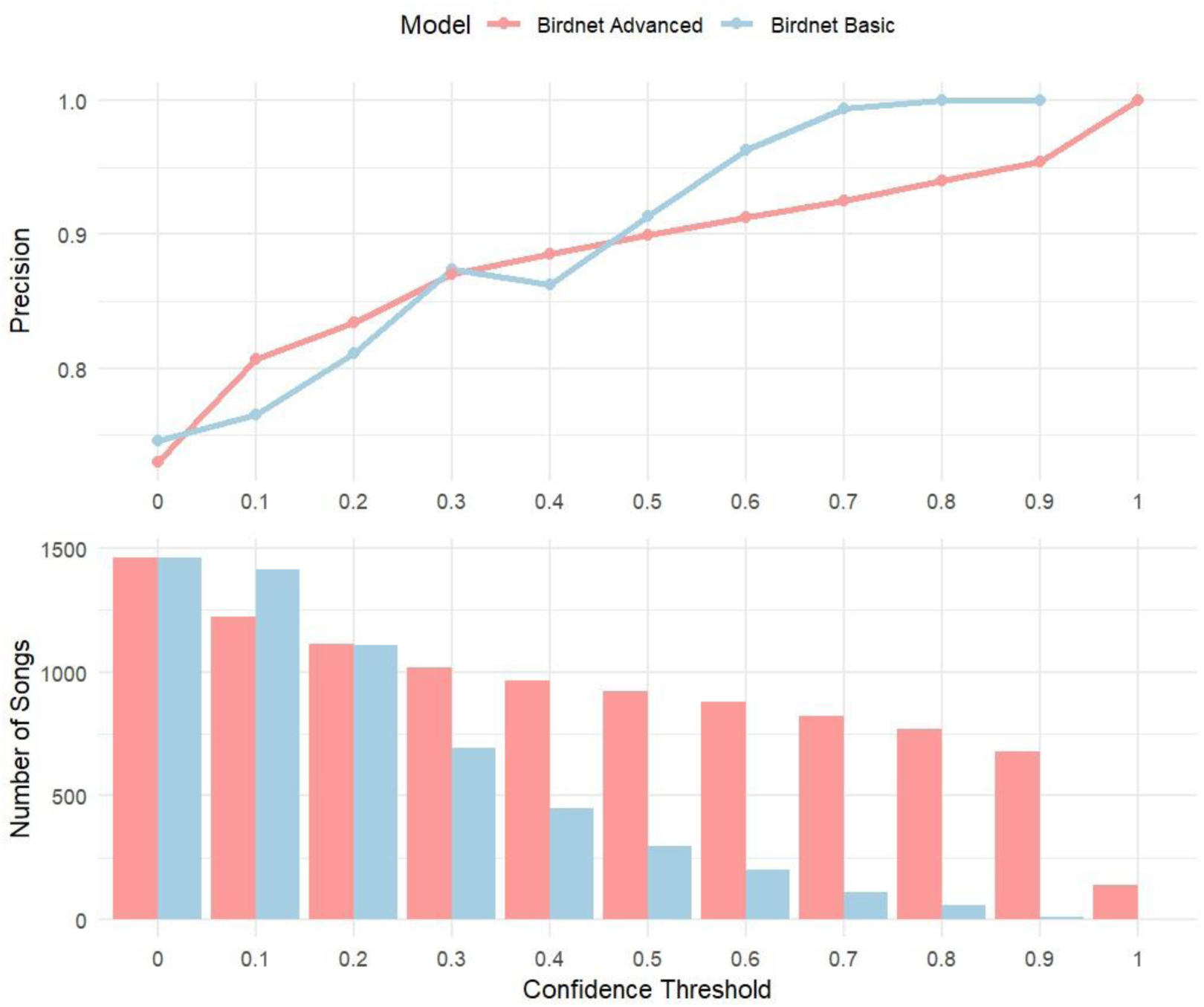
Precision and number of songs retained at different confidence thresholds for *BirdNET Advanced* (red) and *BirdNET Basic* (blue).

### Statistics

The ID detection rate was estimated for the first phrases of Yellowhammer songs at each distance. The number of songs assigned to an individual by each observer was recorded. For the two BirdNET models, a confidence threshold of 0.5 was set to determine whether a song could be assigned to an individual (see above, BirdNET Confidence threshold). Only detections with a confidence score equal to or higher than this threshold were classified as positive. We quantified the ID detection rate as the proportion of calls assigned to an individual at each distance. To account for uncertainty, 95% confidence intervals for these proportions were calculated in R (v. 4.4.2) using the exact binomial test (binom.test). This approach accounts for the discrete nature of detection events and provides an interval estimate for the true detection rate at each distance.

To evaluate the individual identification, the prototypes assigned by the annotator with prior knowledge were considered the ground truth and were compared with those assigned through spectrogram inspections (by humans) and by the two automatic classification models (BirdNET Basic, BirdNET Advanced). The ID assignments were categorized as true positives, false positives or false negatives for each method and distance. Accuracy, precision, and recall were then calculated in a multi-class framework. Accuracy was defined as the proportion of correctly classified cases out the total number of songs in the dataset; Precision as TP/ (TP + FP) and Recall is defined as TP/ (TP + FN), where TP = the number of true positives, FN = false negatives and FP = false positives. We calculated Precision and Recall for each class individually and then computed their macro-average. All performance metrics were computed only for songs consistently assigned to an individual by both human annotators and BirdNET Advanced (confidence threshold of 0.5), thereby ensuring that comparisons were based on the same set of vocalizations and avoid bias from including songs that were considered uncertain ID by either of the observers or models. We did not aim to harmonize the vocalization set to include also BirdNET Basic as this would highly reduce the vocalization set.

We assessed the inter-rater reliability between the two external observers and the BirdNET Advanced model regarding the ID assignment per each distance. The agreement was quantified with the Fleiss’s Kappa (Landis and Koch, 1977) using the formula:

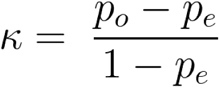

Where *p_o_* is the observed agreement of the raters and *p_e_* is the expected agreement of the raters. The standard error for Fleiss kappa (SE_k_) was calculated according to Cohen (1960) with the following equation calculated:

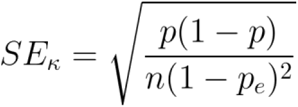

## Results

The degradation of songs seems negligible up to 100 m, where the songs remain still clearly visible in the spectrograms, while from 150 m the degradation substantially reduces the number of discernible elements and alters their temporal and spectral structure, making them increasingly difficult to assign to individuals and sometimes to the species as well. We illustrate the effect of propagation on signal quality on the spectrograms of three types of songs recorded at different distances (Figure 1b). The reduction of the signal-to-noise ratio (SNR) with distance is clearly apparent (see also Supporting information, Figure S2).

### ID detection distance

Human observers reliably detected the simulated Yellowhammer songs from spectrogram inspection at close and middle ranges. All 209 songs were clearly present and assigned to an individual up to 100 m, with an average ID detection rate across human observers of 1.00 (95% CI: 0.98–1) up to 50 m and 0.97 (95% CI: 0.94-0.99) at 100 m. Beyond 100 m, the ID detection rate of the two human observers declined sharply, with substantial variability between them (Figure 3), reflecting the difficulty in assigning identity on fainter calls at longer ranges. ID detection rate for the BirdNET Basic ranged from 0.40 at 6.5 m (95% CI: 0.33–0.47) down to 0.13 at 100 (95% CI: 0.09–0.19). BirdNET Advanced achieved higher ID detection probabilities across distances, from 0.94 at 6.5 m (95% CI: 0.90–0.9) to 0.64 at 100 m (95% CI: 0.57–0.70). Both models showed rapid decline in detection probability already beyond 50 m, at closer distances than in human observers (Figure 3).

**Figure 3.**
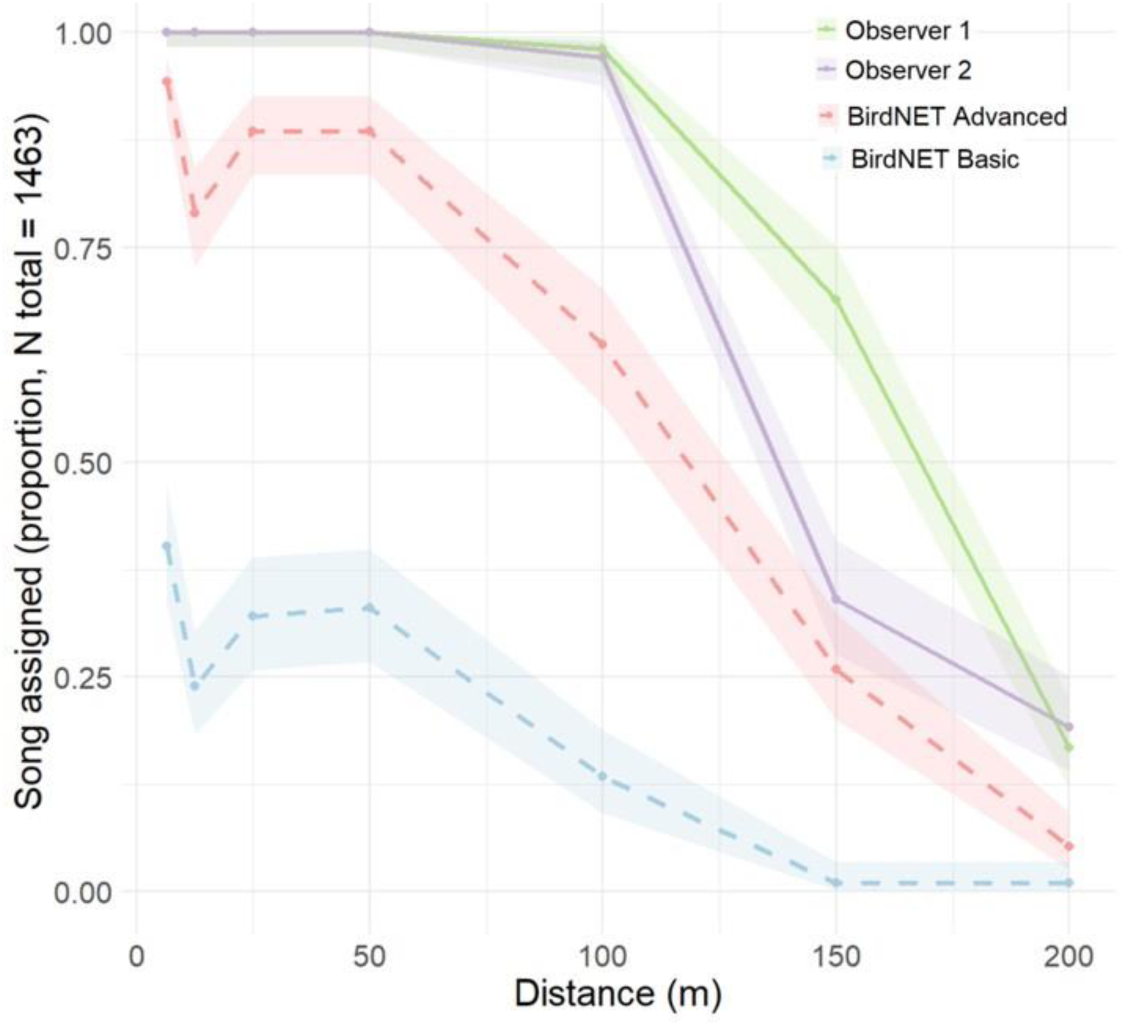
ID detection rate as a function of distance for two automated models (BirdNET Basic and BirdNET Advanced – confidence threshold 0.5) and two human observers. Solid lines represent human observers, while dashed lines represent automated models.

### Success of individual identity assignments

For the analysis of accuracy of individual identity assignments, a reduced dataset was used. Of the 1,463 songs recorded at seven different distances, around 1% (18) were excluded due to overlapping with other sounds, such as songs from different bird species. Additionally, for 16% (238) of the songs, neither of the human annotators assigned an identity. 5% (77) were excluded because identity was assigned by only a single annotator. As expected, the disagreement from spectrogram inspection was for songs recorded at further distances. Furthermore, an additional 21% of songs (241) were excluded by considering only BirdNET Advanced assignments with a confidence score greater than 0.5, resulting in a final dataset of 889 songs in which the identity has been assigned by both human observers and BirdNET Advanced.

The performance of individual identification was evaluated in terms of accuracy, precision, and recall for the two human observers (spectrographic analysis) and the two BirdNET-based automatic models. The performance of the automatic detector decreased substantially with decreasing song quality. All metrics remained consistently high (above 0.80) up to 50 m across all annotators. Starting from 100 m, however, a clear decrease was observed, which was particularly pronounced for the BirdNET Basic, reaching a value of 0.46 in accuracy. From 150 m, both automatic classifiers demonstrated a steep decline in performance, while the human observers maintained higher and more stable values across distances (Figure 4). Considering the performance of the two BirdNET models, the BirdNET Advanced trained with data augmentation outperformed the basic model, achieving results comparable to those obtained from human spectrogram inspections up to 100 m.

**Figure 4.**
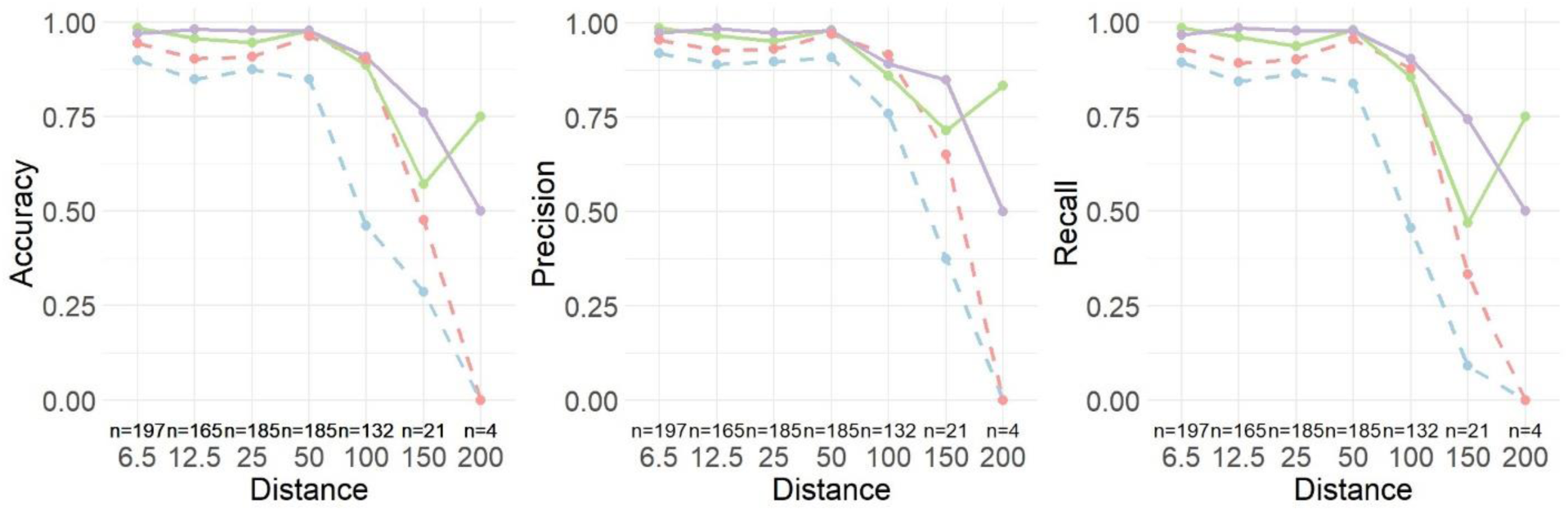
Metrics for individual identification success of models and human observers as a function of distance. From left to right: accuracy, precision, and recall calculated on the reduced dataset (blue = BirdNET Basic, red = BirdNET Advanced, green = Observer 1, violet= Observer 2; solid lines = human observers, dashed lines = automated models).

When we compared the level of agreement among BirdNET Advanced and the two human observers using Fleiss’s kappa statistics, the results showed almost perfect agreement up to 100 metres (Figure 5), and as expected, a marked reduction at longer distances.

**Figure 5.**
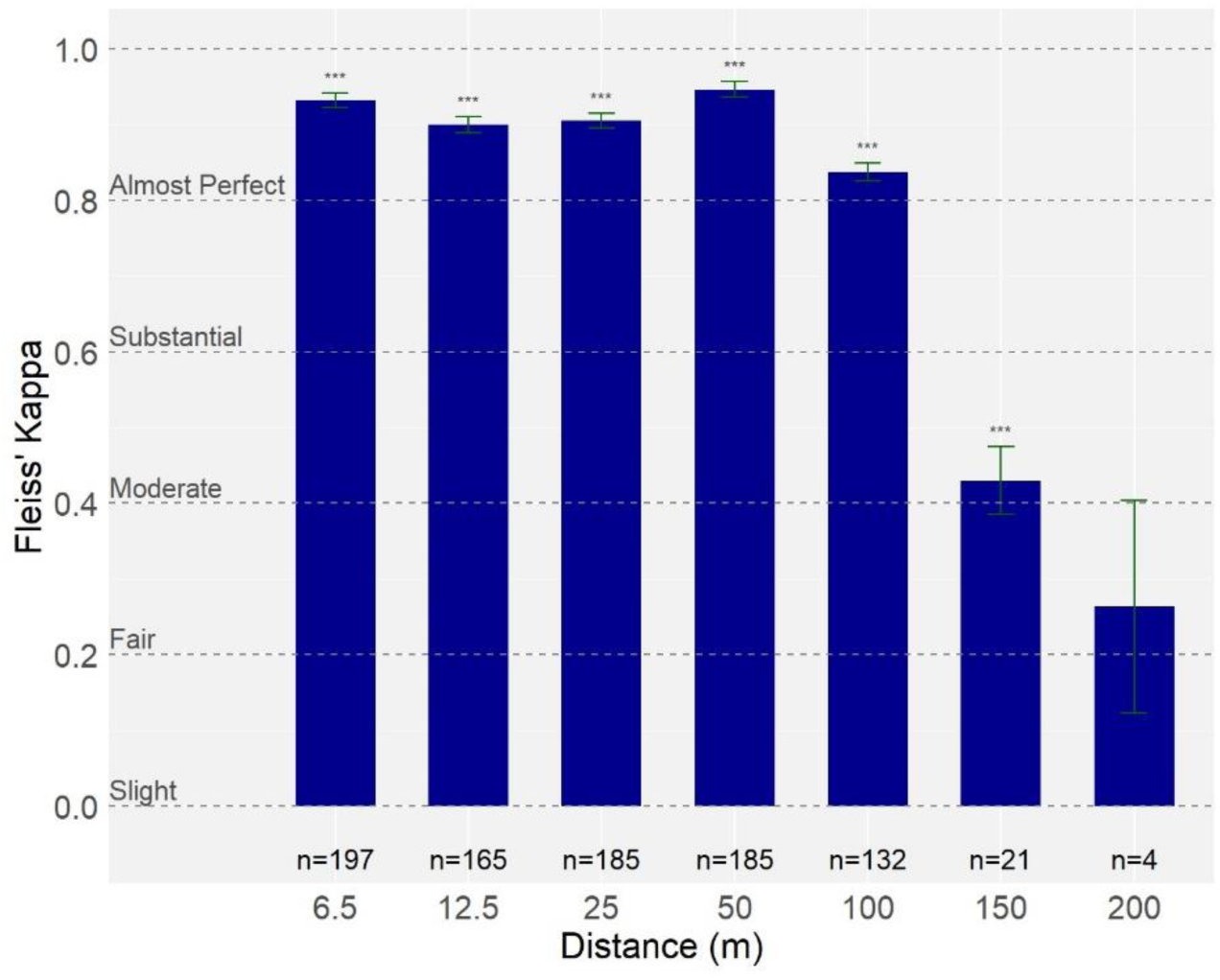
Agreement among the BirdNET Advanced model and the two human observers at different distances, quantified using Fleiss’s kappa statistics.

## Discussion

Our study demonstrates that the possibility to recognize individual Yellowhammers, based on recording of their songs, declines with distance. The majority of broadcasted songs were detectable up to 150 meters, while only a small proportion remained visible in the spectrogram at 200 meters. While Yellowhammer songs could be detected at relatively long ranges, both manual and automatic reliable individual identification was limited to approximately 100 meters. This was manifested in a reduction in the number of songs that could be confidently assigned to a specific individual, and a sharp drop in accuracy of identity assignments after the 100 m category. Relatively simple augmentation tricks boost the performance of BirdNET model to perform almost as good as human experienced observers for assigning song types especially at shorter and moderate distances.

In our study, the detection of Yellowhammer songs extended to 150–200 m, similar to findings reported for other species. For example, Winiarska et al. (2024) reported that small bird species with a frequency range ca. 1,500 and 8,500 Hz typically have a maximum detection distance of around 150 meters. Species’ detection distance directly influences how accurately we can estimate population abundance, species composition, and spatial distribution. Since detection distance is species-specific and affected by habitat (Winiarska et al., 2024) and by the type of recording device used (e.g. Mennill, 2024), it is a key factor in modelling species distributions and assessing conservation status. Knowing how far a species can be detected also helps determine whether it is likely to be recorded or missed and is essential for planning the spatial distribution of ARU deployments (Winiarska et al., 2024). However, the detection alone is not sufficient for individual-based monitoring. Our findings highlight the importance of distinguishing between signal detection distance and individual identity detection distance in acoustic studies. The former refers to the maximum range at which a sound can be detected above background noise, whereas the latter corresponds to the distance at which the acoustic features necessary to discriminate among individuals remain intact, and this distance could be much shorter (100 vs 150m in this study). While songs may be audible over long distances, the range over which they convey individual-specific information is narrower. For example, in the white-browed warbler (*Basileuterus leucoblepharus*), the individual signature encoded in male songs degrades rapidly over short distances, typically less than 100 m, meaning that individual information is largely limited to nearby individuals, such as territorial male neighbors and the female partner. In contrast, species-specific identity is more resistant to degradation and can reach a wider audience, for example to attract other females or deter rival males (Mathevon et al., 2008; see also the little auk single call, Osiecka et al., 2024).

Understanding the distance at which individual identity can still be reliably assigned is critical to move beyond species detection in PAM (Knight et al. 2025). Relying solely on detection distance when deploying ARUs can lead to incomplete coverage of the study area, as the range for reliably identifying individuals is shorter than the maximum detection distance. This may result in missing some individuals, limiting the ability to track movements and producing biased population estimates. Considering individual identification distance ensures that ARU placement captures both the full spatial extent of the population and reliable individual-level data.

In our study, we evaluated both manual and automatic (deep learning-based) approaches to assign individual identity across the increasing recording distances. Automatic individual identification has been explored in several studies and different manual, and also automatic solutions were suggested (see review Knight et al., 2024 but also Zhao et al., 2023; Lapp et al., 2025). Similar to Huang et al. (2025), who used BirdNET for individual-level recognition, we found that pretrained models originally tuned for species classification can also be adapted to discriminate among individuals. BirdNET feature embeddings could successfully differentiate between classes of acoustic events that are very similar and differ only in subtle details (compare e.g. Figure 1a - Song2 and Song3). BirdNET model provided a scalable approach for individual identification. Its user-friendly interface enables users with little or no machine learning experience to process large volumes of audio data rapidly and efficiently. We believe this is a big advantage and a promising step forward for wider adoption of these methods within the community. However, Yellowhammer songs (as well as many other bird species) are particularly suitable to be used with BirdNET model architecture. For example, BirdNET scans recordings using 3s time windows which match well the song durations of Yellowhammers but might not work so well if ID specific features appear on much larger or smaller temporal scales. Other similar BirdNET limitations should be considered for each particular use. Despite the convenience of BirdNET, more flexible computational frameworks, such as transformers (Vaswani et al., 2017) and structured state space sequence models (Gu et al., 2022), might still be preferred for many species with more complex songs (Huang et al., 2025).

One of the main challenges in applying automated tools to passive recordings lies in the low signal-to-noise ratios that often characterize field recordings with individuals vocalizing far from the microphone and prevent effective use of automatic detection methods (Knight et al., 2024). We tested two versions of the BirdNET model: one trained on a restricted dataset containing only songs with a high signal-to-noise ratio (hereafter referred to as BirdNET Basic; mimicking humans getting one good quality example per song type), and another trained with additional data augmentation (hereafter BirdNET Advanced; mimicking some human obvious self-training during the assessment task), which aimed to include distance-related variability into the training by adding noise backgrounds, pitch shifting, and time stretching modifications to original songs. Still, BirdNET Basic performed surprisingly well up to ca. 50 m (Figure 4) despite steadily decreasing SNRs (Supporting information, Figure S2), showing that even the basic model exhibited some level of robustness towards recording quality despite no degraded songs were included in training data. Similarly, Haley et al. (2024) found that BirdNET maintained high detection accuracy when recordings had high signal-to-noise ratios (> 10 dB).

Moreover, relatively simple data augmentation (BirdNET Advanced) significantly enhanced model robustness of individual identification for both close range and middle range distances up to 100 m, almost matching the level of human performance. Humans still performed better for the largest distance categories (150 m and 200 m). It could be possible that our augmentation did not appropriately represent all the degradation processes that occur in real conditions. It would be interesting to see if including real degraded songs from large distances might further improve performance of the model. It also highlights that humans’ cognitive pathways possess particularly good mechanisms for processing noisy signals even when confronted with new tasks without available training data. Nevertheless, our results confirm that model fine-tuning and data augmentation are essential steps to enhance generalization and reduce overfitting, especially when working with a limited number of samples available for training, a common limitation in the bioacoustics field (Knight et al., 2024).

Another critical aspect when applying BirdNET Analyzer to large acoustic datasets is the selection of an appropriate confidence score threshold (Wood et al., 2023; Amorós-Ausina et al., 2024; Singer et al., 2024). Similarly to Pérez-Granados (2025) who tested the performance of BirdNET Analyzer at different distances, we found that the number of songs with high confidence scores declined significantly with increasing distance from the source. Setting up lower confidence thresholds increases detection rates but may result in false positives (FP), while higher thresholds improve reliability at the cost of excluding valid detections. The trade-off between precision and detection range can complicate bird richness and density estimations if the detection radius becomes overly limited (Pérez-Granados, 2025) or detections become unreliable. These problems associated with species detection ranges also arise with individual identity detection ranges.

We studied the possibility of AIID from a human perspective only. It remains to be tested how far conspecifics themselves could recognize other individuals. It has been shown that male Yellowhammers use songs in territorial disputes (Hiett and Catchpole, 1982) and to discriminate between neighbors and strangers, which might help them to reduce unnecessary territorial conflicts (Hansen, 1984). It is not known which features Yellowhammers use to identify individual conspecifics. Their songs are composed of trills (the initial phrase) and whistles (the dialect phrase). Previous studies indicate that whistles transmit more effectively over long distances than broadband trills (Wiley and Richards, 1982; Slabbekoorn et al., 2002; Naguib, 2003) and in Yellowhammers’ individual identity information, at least from human perspective, seems to be encoded in the fine structure of syllables in the broad band trills of the initial phrase that we focused on in this study. Birds typically express better temporal and frequency resolution of hearing compared to humans and are able to perceive small details in the syllable structure (Dooling et al., 2002; Dooling and Prior, 2017).

However, both manual and automatic individual identification approaches in this study were based on spectrograms with sufficient resolution making it possible to use tiny time and frequency-modulation details for individual identification. Deterioration of spectro-temporal information in these elements would be particularly critical for individual recognition at increasing distances. High-frequency song elements in Yellowhammer songs are especially susceptible to attenuation and distortion in open habitats. This pattern aligns with findings in other passerine species, where spectral features degrade faster than temporal ones during transmission, leading to a more rapid loss of identity information than of detectability (e.g., White-browed Warbler: Mathevon et al., 2008; Zebra Finch: Mouterde et al., 2014). It would be interesting to investigate the features that Yellowhammers themselves use for individual recognition and whether final phrase whistles or, for example, temporal individual information (Osiecka et al., 2025) could function as complementary or even more robust components of individual identity signaling. However, it is interesting to note that our threshold for individual identification (ca. 100 m) matches well with the reported size of territories, and the distances over which males typically interact. Most foraging trips occur within 100 m of the nest (Morris et al., 2001), and Yellowhammers spend the majority of their time within this range (Biber, 1993; Morris et al., 2001; Anderson, 2014). This suggests that the individual identity detection distance reported in this study, indeed, might be ecologically relevant and delineate a space where Yellowhammers can efficiently defend their territories and manage social relationships with their neighbors.

### Dataset potential and limitations

This study and the associated dataset were intentionally designed to evaluate how detection models perform under varying recording quality, particularly at low signal-to-noise ratios. Recording acoustic signals under different environmental conditions and levels of degradations represent a well-established principle on how to make recognition models more robust (e.g. Kahl et al., 2021; Ghani et al., 2025) and perform well under real-world conditions. Still, this approach has been mostly neglected in acoustic studies and similar datasets with well-defined conditions (e.g. distance from source) are in general not available (Gibbons et al., 2025; Weldy et al., 2025). Understanding how acoustic signals vary with distance, recording devices, and species can provide valuable insights for bioacoustics monitoring. Such knowledge can help define the effective sampling area of passive acoustic monitoring (PAM) surveys. By incorporating distance-related variability into model training data, it becomes possible to enhance model robustness and better generalize to real-world conditions. Our dataset was already used to develop lightweight and efficient models for on-device and sustainable bioacoustics monitoring (Carmantini et al., 2025). Datasets similar to ours can also be used to develop algorithms that could better infer and predict how degradation might affect high quality samples provided for training similar to how humans tackle this problem. Therefore, it may contribute to broader ecological applications, including tracking individual animals, developing site occupancy models, and estimating bird richness or density around recording devices.

The dataset used in this study consists of recordings from only ten individuals limiting the generalizability of the results. Further, the present experiment represents a closed-set classification scenario, where both training and test datasets include the same individuals. In these conditions, the model performed well, demonstrating its potential for applications in contexts where the local population is well known, for example, long-term monitoring projects where a catalog of individuals is available and the territory is well characterized. This approach may be particularly well-suited for geographically constrained populations, such as island species or isolated breeding colonies, where it is methodologically feasible to record most individuals within the population. Further, recent study has shown that pre-trained models fine-tuned on a subset of known individuals can be extended to handle previously unknown individuals (Huang et al., 2025). It is also important to note that the recordings used in our study were collected under relatively favorable conditions (i.e., low background noise and a loudspeaker facing the recorder), which likely contributed to the consistent recognition performance observed at close and intermediate distances. In natural environments, multiple factors may affect the effective range of species and individual recognition, including habitat structure (Yip et al., 2017), ambient noise levels, weather conditions and or microphone quality (Darras et al., 2020). Additionally, overlapping vocalizations, due to either the interactions between individuals of the same species (counter-singing) or the presence of other species, may further complicate individual identification. Future works should therefore evaluate the robustness of our approach across diverse acoustic environments, recording devices and ecological contexts.

## Conclusion

Overall, our findings illustrate both the potential and the current limitations of individual identification using low-cost passive recording devices and ready-to-use machine learning tools. Promising tools for large-scale individual monitoring are available. Their effectiveness ultimately depends on careful dataset preparation, threshold optimization, and validation against expert-based annotations. Moving forward, progress in this field would benefit from integrating standardized transmission experiments into the design of passive acoustic monitoring networks, to guide the optimal deployment of autonomous recorders and to provide valuable data for fine-tuning automatic detection algorithms. Including recordings from transmission experiments enable diversification of training datasets to capture recording distance effects and habitat-specific acoustic properties. Such an integrated approach will be essential to enhance robustness of detection models and ensure their applicability to real-world monitoring scenarios, ultimately enabling reliable and scalable bioacoustic tools for ecological research and conservation.

