## Supporting information, Figure S1 for "Assessing the Effective Range for Individual Acoustic Identification: Comparison of Manual and Automatic Methods"

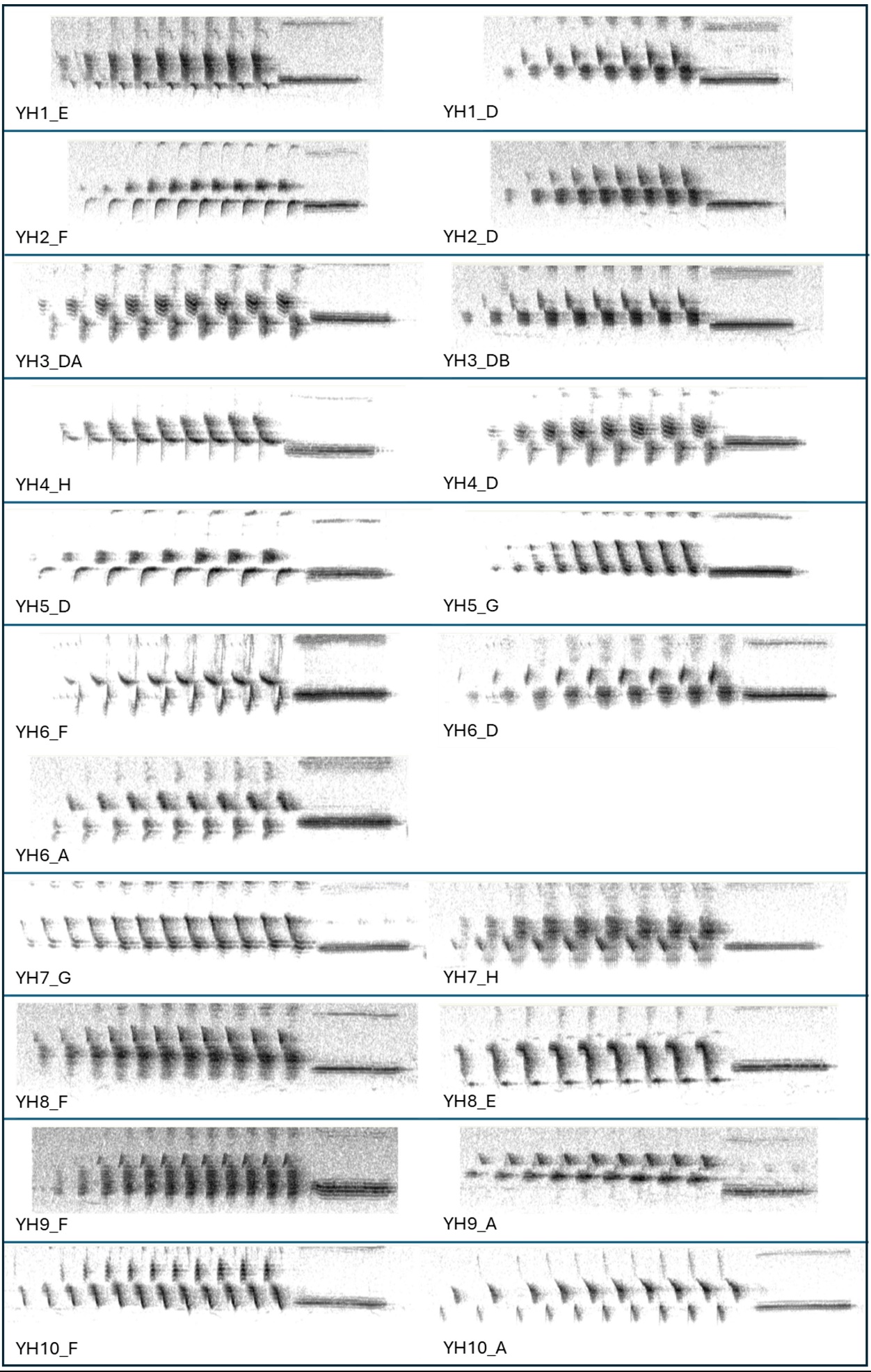

Figure S1

Catalog of the 10 male Yellowhammers (*Emberiza citrinella*) used in the transmission experiment. Each line represents the spectrogram (frequency range: 3000 to 8000 Hz) of a different male and includes its entire repertoire of initial phrase types used as playback stimuli. Each prototype is defined as a combination of a broader song phrase type category (A–H) and an individual variant of that phrase type corresponding to the male ID (1–10).
