## Supporting information, Figure S2 for "Assessing the Effective Range for Individual Acoustic Identification: Comparison of Manual and Automatic Methods"

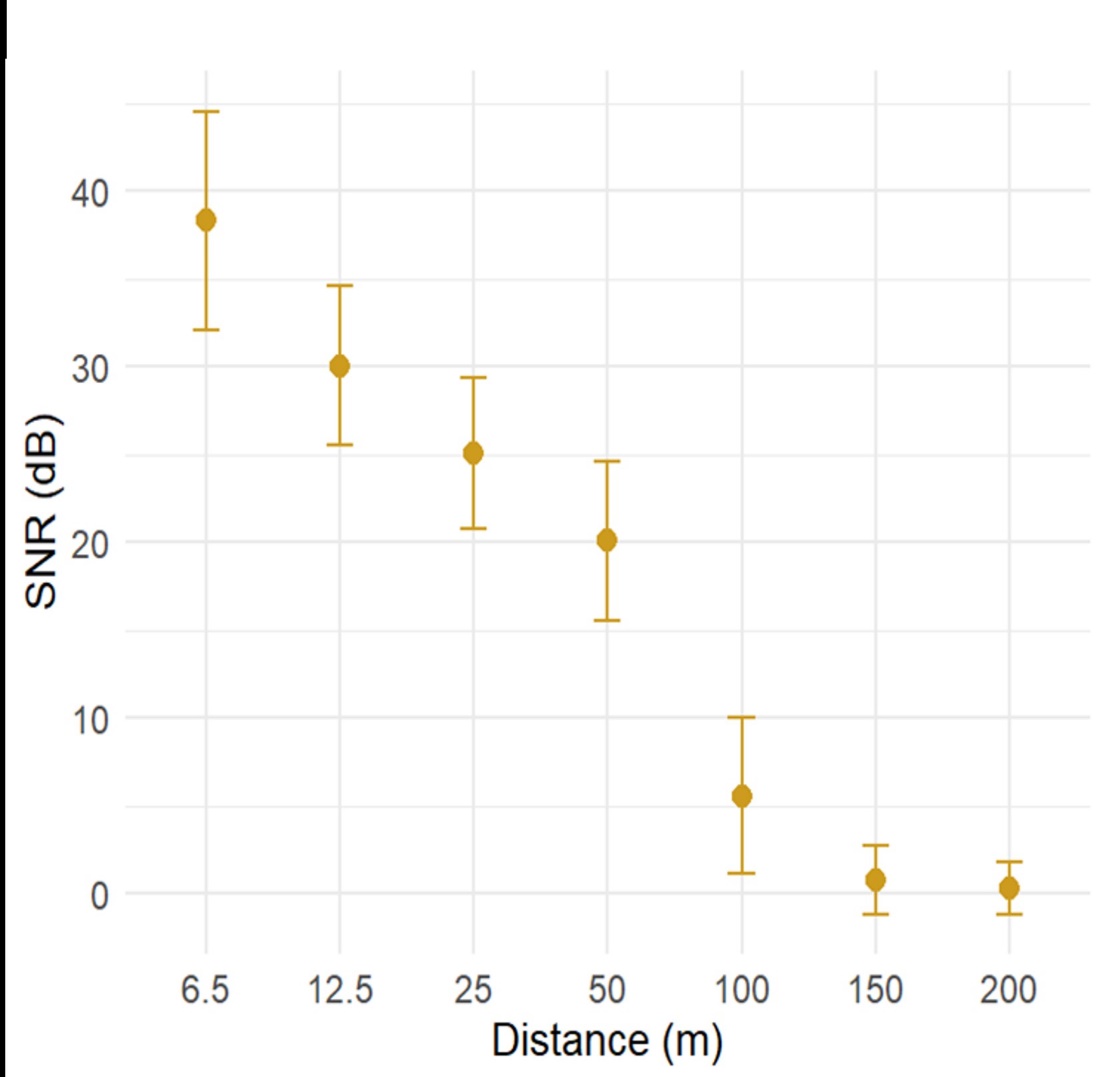

Figure S2. Plot of the signal-to-noise ratio level as a function of the source-to-receiver distance. For each distance, the distribution of SNR values is represented through their main statistics: average, minimum and maximum.
