## Supporting information, Table S1 for "Assessing the Effective Range for Individual Acoustic Identification: Comparison of Manual and Automatic Methods"

Table S1. Metrics for each BirdNET model across different confidence thresholds varying from 0 to 1, including the support (number of retained songs), accuracy, precision, recall, and F1 score.

|  | metric | 0 | 0.1 | 0.2 | 0.3 | 0.4 | 0.5 | 0.6 | 0.7 | 0.8 | 0.9 | 1 |
| --- | --- | --- | --- | --- | --- | --- | --- | --- | --- | --- | --- | --- |
| BirdNET Basic | Support | 1463 | 1415 | 1107 | 692 | 449 | 298 | 199 | 108 | 55 | 9 | 0 |
|  | Accuracy | 0.56 | 0.57 | 0.62 | 0.77 | 0.90 | 0.96 | 0.98 | 0.99 | 1.00 | 1.00 | Na |
|  | Precision | 0.74 | 0.76 | 0.81 | 0.87 | 0.86 | 0.91 | 0.96 | 0.99 | 1.00 | 1.00 | Na |
|  | Recall | 0.55 | 0.56 | 0.60 | 0.73 | 0.85 | 0.90 | 0.97 | 1.00 | 1.00 | 1.00 | Na |
|  | F1 score | 0.59 | 0.60 | 0.50 | 0.76 | 0.84 | 0.90 | 0.96 | 0.99 | 1.00 | 1.00 | Na |
| BirdNET Advanced | Support | 1463 | 1223 | 1112 | 1015 | 965 | 920 | 878 | 820 | 767 | 680 | 139 |
|  | Accuracy | 0.65 | 0.75 | 0.80 | 0.84 | 0.87 | 0.89 | 0.90 | 0.92 | 0.94 | 0.96 | 1.00 |
|  | Precision | 0.73 | 0.81 | 0.83 | 0.87 | 0.88 | 0.90 | 0.91 | 0.92 | 0.94 | 0.95 | 1.00 |
|  | Recall | 0.65 | 0.79 | 0.79 | 0.84 | 0.86 | 0.88 | 0.89 | 0.91 | 0.92 | 0.93 | 1.00 |
|  | F1 score | 0.67 | 0.76 | 0.80 | 0.84 | 0.86 | 0.88 | 0.89 | 0.91 | 0.92 | 0.94 | 1.00 |
